# Gene network organization, mutation and selection collectively drive developmental pattern evolvability and predictability

**DOI:** 10.1101/2024.12.23.630099

**Authors:** Harry Booth, Z. Hadjivasiliou

## Abstract

A hallmark of development is the generation of spatial patterns driven by morphogen gradients and Gene Regulatory Networks (GRNs). Although the mechanistic basis by which GRNs orchestrate cellular responses and tissue patterning during development is well understood, their evolutionary dynamics remain less clear. Still, utations in regulatory elements that govern GRN-driven patterning are a key mechanism for patterning evolution. In this study, we use the *de novo* evolution of a stripe phenotype as a model framework to investigate the evolutionary dynamics associated with the emergence of new spatial gene expression boundaries and the adjustment of existing boundaries. To probe general principles of GRN-driven pattern evolution we introduce a new high-throughput theoretical framework that rapidly produces a comprehensive dataset of evolutionary trajectories. We leverage this large dataset to investigate the types of mutations that drive different phenotypic shifts in spatial patterning. Our findings suggest that the order in which mutations in gene-gene interactions appear, and the specific combination of gene-gene interactions that mutate together determine the evolvability of novel spatial gene expression patterns. We interpret our results in the context of epistatic effects that naturally arise in networks of interconnected genes, and show how contingencies and constraints emerge in our system. Our results elucidate the interplay between mutation and gene network organization, revealing how historical contingencies arise and impact the evolvability of GRNs and the predictability of their evolutionary outcomes.

## Introduction

A central challenge in evolutionary developmental biology is to elucidate how phenotypic variation arises from underlying developmental mechanisms. At the core of development lies the spatial patterning of tissues into different cell types, driven by gene regulatory networks (GRNs) that interpret morphogen gradients^1^. While the mechanistic basis of how GRNs influence spatial patterning during development is well-established, the interplay of these networks with evolutionary forces such as mutation, selection, and historical contingency remains poorly understood. Yet, several studies have revealed the critical role of GRNs in generating morphological variation both within and across species^2–6^, underscoring the importance of investigating their evolutionary dynamics in this developmental context.

Understanding how mutations at the sequence level impact GRN function and their evolutionary dynamics is a key milestone for the study of GRN-driven pattern evolution. The current consensus is that GRN evolution advances through mutations not in the regulatory genes or transcription factors (TFs) themselves, but in cis-regulatory elements (CREs) or enhancers that drive TF expression levels^6–8^. In this way pleiotropic effects associated with TF mutations are bypassed, while mutations in regulatory regions are able to generate variation for selection to act upon. Therefore, the ways in which mutations in CREs reorganize gene interactions and the associated GRN topology is central in understanding the evolution of GRNs and the spatial patterns they coordinate during development.

Recent work has investigated the role of point mutations in native CREs and the insertion of *de novo* CREs into a GRN controlling the patterning of fruit fly embryos. This revealed that the impact of mutations on the phenotype depends on whether mutations affect existing CREs or introduce new regulatory elements in the system^9^. New assays that allow targeted genetic manipulation of GRNs *in vivo* can generate large datasets of mutations at the regulatory and TF level linked to phenotypic shifts^10,11^. Furthermore, synthetic approaches to patterning network evolution offer promising tools for investigating the mechanistic role of genes, their interactions in patterning networks and impact on evolution^12,13^.

However, it is currently unfeasible to characterize the GRN re-organization which may result from high throughput experimental mutation assays at a scale large enough to extract common principles of network re-wiring in these systems. Furthermore, the difficulty of imposing a selection pressure for novel patterns in multicellular development means that the role of this network re-wiring within an evolutionary process is challenging to analyze. Hence, although new experimental tools promise to advance our understanding of the evolution of development, the complexity of GRN-driven patterning evolution, characterized by multi-scale, multi-component, and non-linear dynamics, coupled with the timescales over which evolutionary change occurs, present challenges that experimental tools alone may struggle to overcome. Since mutation arises at the sequence level, a comprehensive understanding how mutation drives network re-wiring must account for the complexity of the sequence-to-network mapping. However, this bottom-up approach to understanding patterning evolution can be complemented with a top down approach that focuses on genes and their network of interactions. More minimal, mechanistic models are tractable while having the potential to capture physical constraints emerging from the collective transcriptional dynamics, which are relevant in the evolution and diversification of patterning mechanisms. This approach requires a mechanistic description of GRN-driven patterning integrated with mutation and selection, and forms much of the motivation behind this work.

A related question and long standing debate in evolutionary biology is the relative role of stochasticity and constraints in determining evolutionary outcomes. Historical contingency during evolution describes a sensitivity to past events, that are often inconsequential at the time they occur, but that nonetheless impact subsequent evolution. This sensitivity becomes heightened when adaptation takes place on “rugged” fitness landscapes^14^, a property which emerges through the complex, non-linear nature of many biological genotype-phenotype mappings^15^. Whilst historical contingency has been investigated using evolutionary simulations in protein evolution^16^, and in GRNs with simple evolved functions^17^, these investigations have yet to be extended to genotype-phenotype maps defined by multicellular development processes such as patterning. For example, whilst it is known that spatial pattern accessibility is dependent on GRN topology^13,18^, we do not yet understand at what point in the evolution of new developmental patterns network constraints might become influential, or how these constraints interact with other evolutionary forces. For instance, neutral evolution - whereby mutations with little or no impact on the phenotype can accumulate - has been proposed as a potentially important force in GRN-driven pattern evolution^12^. But the ways in which nearly neutral mutations may impact future evolution and diversification of GRN topology and spatial patterns is not understood. Theoretically, multiple GRN topologies can drive the same spatial pattern at the tissue scale^19^, which poses the question how and why certain patterning mechanisms may have evolved in preference to others.

Theoretical models of GRN-driven patterning have successfully captured the spatiotemporal patterning dynamics observed in various model systems, from the *Drosophila* embryo^20–22^ to the vertebrate neural tube^23,24^. While these models do not explicitly track genetic information at a molecular scale, they have provided crucial mechanistic insights into the fundamental characteristics and emergent properties of GRN patterning^25–27^, and offer a promising and tractable framework for exploring GRN-driven pattern evolution. Previous work integrating mechanistic GRN patterning models into evolutionary simulations has yielded valuable insights into the constraints and design principles that optimize specific phenotypic outcomes, or “fitness”^28–31^. For example, simulations have identified parsimonious paths in GRN space linking Dipteran species and their homologous relationships^32^, elucidated the design principles underlying body plan segmentation in flies^33^, and explored potential routes toward the *de novo* evolution of body segmentation^34,35^. This mechanism-centered approach highlights the central role of gene interactions and network topology – rather than individual genes – as key drivers of evolutionary processes. A common limitation of these approaches to date has been the efficiency at which they can generate evolutionary trajectories. As a result, previous studies have been limited to datasets of evolutionary repeats that were not large enough to investigate general principles of mutation and selection in a developmental context.

In this work we explore the interplay between mutation, gene network organization and natural selection in shaping the evolution of GRNs and developmental patterns. We specifically use stripe-forming networks as a framework to investigate how new gene expression boundaries are formed and how existing gene expression boundaries diversify. We present a high-throughput theoretical framework that integrates a mechanistic description of GRN patterning with evolutionary parameters. By leveraging the framework’s capacity to rapidly evolve target phenotypes, we produce an extensive dataset of independent evolutionary trajectories. This allows us to employ statistical tools to investigate general principles that emerge in our evolution experiments. By employing computational and machine learning techniques we 1) investigate the types of mutations that establish *de novo* gene expression boundaries in space, or refine existing boundaries and 2) build a predictive framework of GRN pattern evolution that we use to identify sources of contingency and convergence as gene networks evolve and complexify. In this way we were able to identify how constraints that emerge from the mutational process and current topology of a patterning GRN may impact developmental pattern evolvability. Our framework offers a powerful and adaptable platform for the investigation of GRN-driven pattern evolution, that has the potential to link physical constraints that emerge from genetic architecture to evolutionary change and diversification.

## Methods

### A high-throughput framework for GRN evolution

We use the *de novo* evolution of a stripe as a model framework to investigate the evolutionary dynamics associated with the emergence of new spatial gene expression boundaries and the adjustment of existing boundaries. We consider a minimal GRN downstream of a morphogen and introduce mutation of the regulatory interactions between genes. Our theoretical framework encompasses three key components (fig. 1): i) GRN-mediated spatial patterning, ii) mutation and fitness evaluation, and iii) a Monte-Carlo scheme which simulates an evolutionary trajectory.

**Figure 1:**
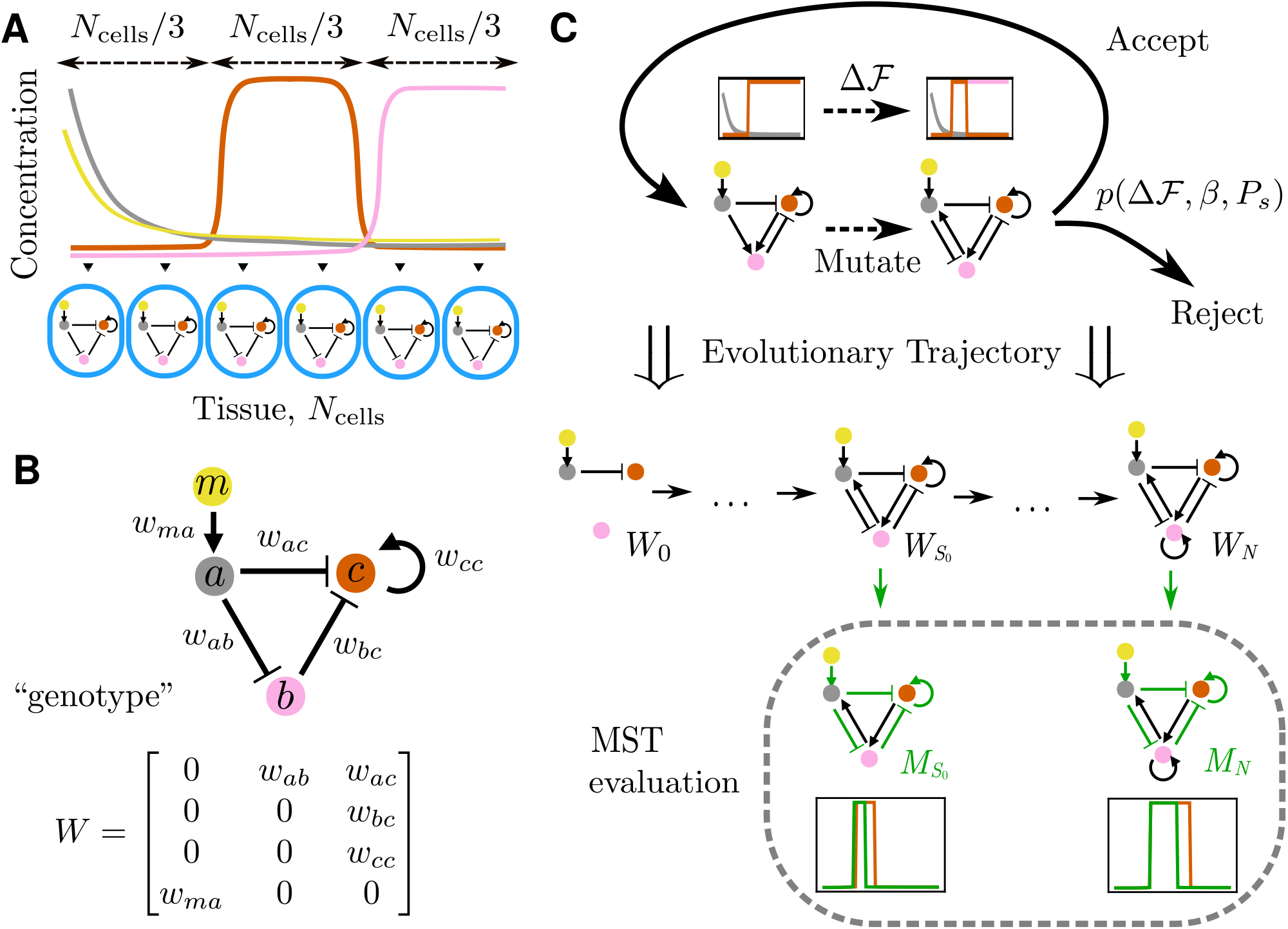
Method Summary. A. Model of developmental patterning and target stripe. A tissue is made up of a discrete number of cells (blue ovals), each with an identical 3-node gene regulatory network (GRN). A morphogen gradient (yellow) is assumed to be at steady state across the tissue. The target stripe profile (orange) has boundaries which define three equal size regions across the tissue. The gray and pink circles represent the two other genes in the network. Lines in the plot are a schematic of the putative concentration at steady state for the morphogen (yellow) and three genes shown in the diagram below (gray, orange, pink) B. An example GRN. The morphogen, *m* (yellow) activates gene *a* (gray), which then regulates the output gene *c* (orange) directly and through an intermediary gene *b* (pink). The regulatory organization is specified by a matrix *W* that holds the weights that define an effective strength for activation (positive weights) and inhibition (negative weights) arrows, and represents the genotype in our analysis. C. A schematic outlining the Monte-Carlo scheme for simulating evolutionary trajectories. The resident genotype associated with the current phenotype is mutated by introducing or editing existing interactions (weight changes). The new phenotype defines a fitness differential Δℱ, which then determines the probability *p*(Δℱ, *β, P*_*s*_) of fixation for the mutant based on the selection strength *β* and population size *P*_*s*_. This process then repeats until a predefined fitness threshold is reached. The sequence of accepted mutants is collected to form an “evolutionary trajectory”. The MST (*minimal stripe topology*) associated with the first stripe forming mutant 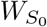 and the final mutant *W*_*N*_ are then evaluated. The MST is the topology associated with the minimal set of regulatory interactions able to maintain a stripe phenotype. Details of parameters used in the evolutionary algorithm are provided in SM.

We consider a 1-dimensional tissue composed of a discrete number of cells, over which a morphogen gradient is established (fig. 1.A). Each cell has an identical GRN consisting of 3 genes {*a, b, c*}, with gene *a* activated by the morphogen, *m* (fig. 1.A-B). The network is specified by a weight matrix *W* whose entries *w*_*ij*_ encode the pairwise regulatory interactions between genes (fig. 1.B). We then assume that the concentration dynamics of gene product *j* in cell *i* ∈ {1, …, *N*_cells_} satisfy

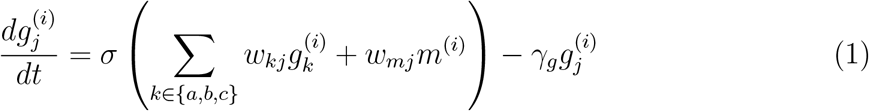

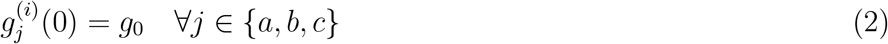

where 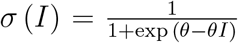 is a sigmoidal activation function with corresponding steepness parameter *θ >* 0 and *γ*_*g*_ is the gene product degradation rate. This description of gene regulation is widely used in both theoretical and experimental studies of spatial patterning in biological systems^19,22,36^.

The evolutionary starting point in our simulations is given by an initial weight matrix, *W*_0_, with a minimal number of non-zero interactions (fig. 1.C), representing an initial state for network evolution. Subsequent mutation adds new regulatory interactions between genes, or adjusts the magnitude of existing ones. Thus, the weight matrix *W* represents the genotype in our model, and mutation acts directly on its entries to produce a mutant GRN.

Previous work has suggested that multiplicative weight changes most accurately represent the effect of point mutations on binding affinities in transcription factor (TF) binding sites^37^. We model this as 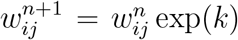, where *k* is normally distributed with 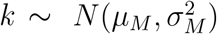 and 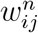 and 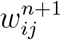 represent the regulatory effect of gene *i* on *j* before and after mutation respectively. Typically point mutations in binding sites decrease affinity^37,38^, and so we set *µ*_*M*_ *<* 0. Whilst point mutations can generate *de novo* regulatory interactions^39^, GRN evolution is thought to occur more efficiently though alternative mechanisms such as duplication^40,41^ or co-option^6,8^. Therefore, to enable mutations of larger effects we also consider additive weight changes such that 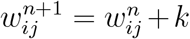, where *k* is normally distributed with 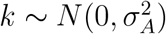. This allows the distribution of weight change magnitudes 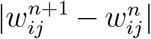 to be independent of the initial magnitude 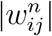. In particular additive mutations enable the introduction of novel interactions (i.e. 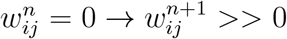). Furthermore, additive mutations enable the sign of a regulatory interaction to change from activating to inhibiting or vice versa.

To investigate the impact of mutations that change a single versus multiple gene interactions in the network, we assume that a single mutant may carry multiple weight changes simultaneously. We implement this by allowing each weight in *W* to change independently with probability *p*. This generates a distribution of weight changes per mutant dictated by *p* (SM, Section 1.2.1). Note that *p* could be interpreted to reflect constraints which emerge from the architecture of CREs and TF interactions, and is hence separate from the mutation rate. For instance, multiple weight changes could occur simultaneously when single mutations on the same CRE affect the binding affinity of multiple TFs, or when TFs themselves mutate. In this way, our framework is set up to investigate how constraints emerging from genetic architecture may impact evolutionary trajectories and evolvability. We return to the implications and biological interpretation of this assumption in the results section and the discussion.

We define a target stripe by a specific width and position (fig. 1.A), and compute the fitness differential Δℱ of mutants relative to the resident phenotype (fig. 1.C). The phenotype *ϕ* upon which selection acts is taken to be the steady-state expression of gene *c* (fig. 1.A). Fitness is evaluated in a two-stage process employing two distinct fitness functions - ℱ_*S*_ and ℱ_*R*_. ℱ_*S*_ is an indicator function which recognizes the transition to a stripe by evaluating phenotypes against a set of general stripe criteria (see SM, Section 1.3). These conditions ensure a stripe-like pattern, but they do not enforce any particular boundary locations. ℱ_*R*_ measures the average distance of the steady state expression of gene *c* to a target stripe pattern with specific boundaries. The target pattern is a stripe that splits the tissue into three equally sized parts (fig. 1.A). Selection for this phenotype can be imposed through the following fitness functional:

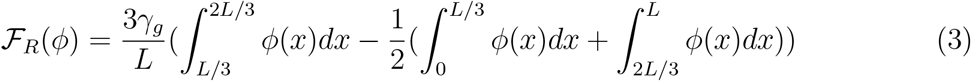

where *L* is the tissue length and *γ*_*g*_ is the gene product degradation rate. The first integral rewards high expression of the gene product in the central region of the tissue. This is then penalized by the average expression of the gene product in the neighboring regions. Finally, this is normalized into the range [−1, 1] by dividing by the upper bound on the difference between the first and second terms. The fitness differential Δℱ between the resident phenotype *ϕ* and mutant phenotype *ϕ*^*′*^ is then evaluated as ℱ_*S*_(*ϕ*^*′*^) − ℱ_*S*_(*ϕ*) if ℱ_*S*_(*ϕ*^*′*^) ≠ ℱ_*S*_(*ϕ*) and ℱ_*R*_(*ϕ*^*′*^) − ℱ_*R*_(*ϕ*) otherwise. This choice is motivated by the fact a phenotype may display a stripe-like pattern whilst not necessarily having a close match in expression space to a stripe with specific boundaries. In these instances we assume its fitness to be higher than a phenotype with no stripe-like characteristics.

This combination of selection functions is designed to generate conditions that favor: 1) the generation of new gene expression boundaries and 2) the adjustment of the position of existing boundaries. In this way our evolutionary algorithm is set up to investigate the network re-organization, and associated mutations, necessary to evolve novel gene expression boundaries and adjust existing ones. The formation and adjustment of gene expression boundaries is a hallmark of developmental patterning and its evolution^1,42^ and so this approach has the potential to yield general insights about the evolution of development.

We model monomorphic evolutionary dynamics with a sequential fixation model, by assuming that the rate at which mutants appear is lower than the time it takes for them to fix^43–45^. The Monte-Carlo scheme for an evolutionary trajectory is defined by the probability that a mutant fixes in a population of size *P*_*s*_:

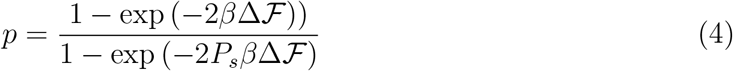

where *β* represents the strength of selection^46^. Refer to SM, Section 1 for further details on the dynamical equations, evolutionary algorithm and parameters used.

## 1 Results

### Multiple GRN topologies evolve following selection for stripe phenotype

We use the algorithm defined in the previous section to generate ~ 10^5^ independent evolutionary trajectories from the initial conditions to the target phenotype. We start our analysis by developing a method to classify the stripe-forming mechanisms associated with networks that evolved across our simulations. To do this, we recorded the evolutionary trajectory 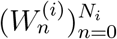 of accepted genotypes, where *N*_*i*_ is the total number of accepted mutants leading to the target phenotype for the *i*^*th*^ trajectory (fig. 1.C). We found that 88.2% of our simulations converged to the target phenotype within 250,000 generations (SM, Section 4, Figure M8.A). Across evolutionary trajectories we defined four key stages; the first accepted mutant (*n* = 1), a period of pre-stripe formation for *n* ∈ (1, *S*_0_), stripe inception at *n* = *S*_0_ and post-stripe formation for *n* ∈ (*S*_0_, *N*_*i*_]. Therefore, *S*_0_ is the mutant number for which a stripe first manifests (fig. 2.A).

**Figure 2:**
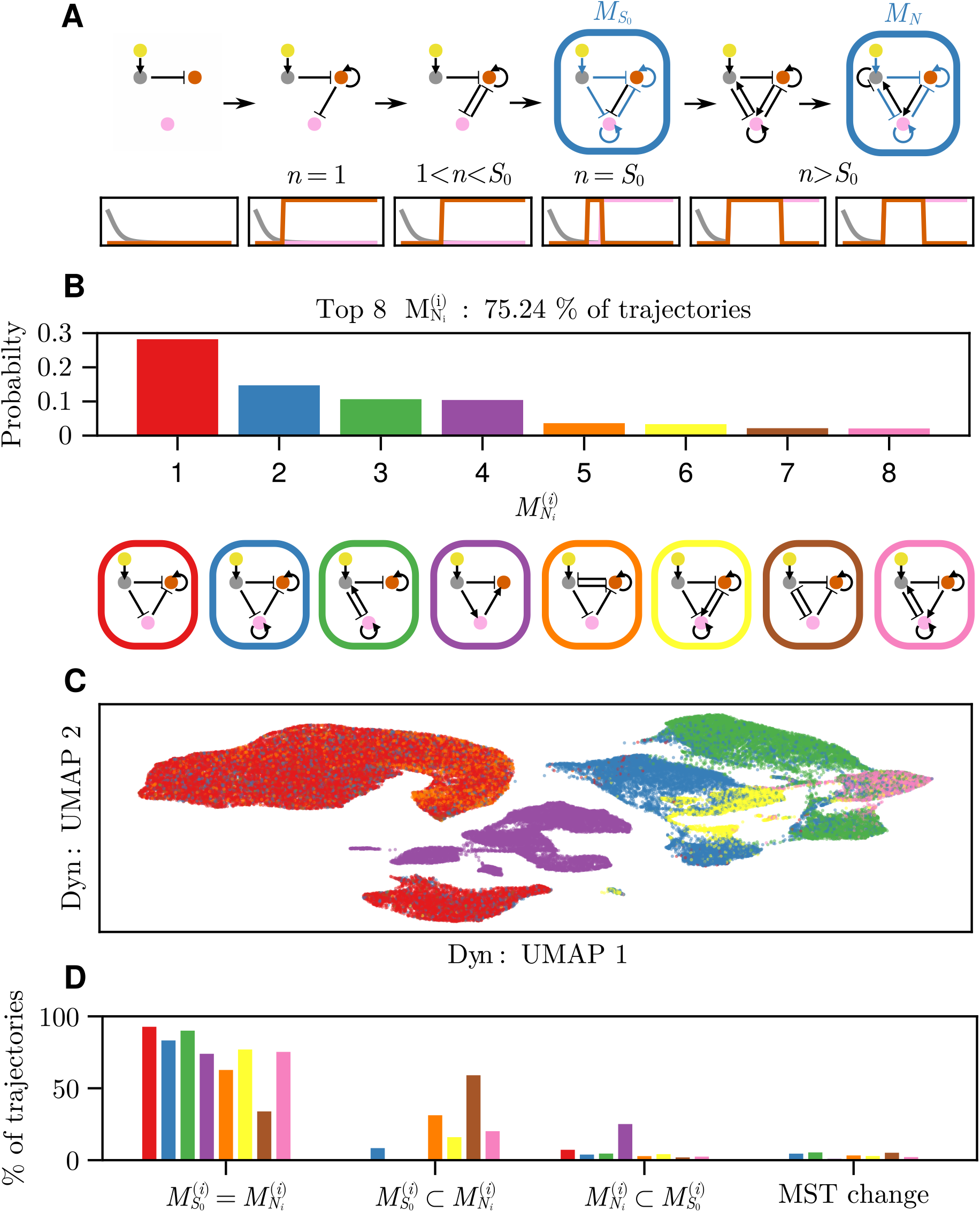
The distribution of adaptive outcomes relating to the core mechanisms of stripe formation. A. An example evolutionary trajectory showing topological changes in the underlying network along with the associated phenotypes. Each step indicates an accepted mutant. The change in the network topology (genotype) is shown on top with the corresponding gene expression profile (phenotype) below. The value of *n* is the index of the shown accepted mutant. The interactions which form the MST at *n* = *S*_0_ and *n* = *N* are highlighted and denoted as 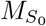 and *M*_*N*_ respectively. B. The distribution of top 8 MST outcomes 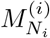 across evolutionary simulations. Colors correspond to the MSTs visualized below the distribution. C. A UMAP^85^ embedding of the dynamics associated with final evolved networks 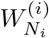, with color coding indicating MST classification shown in B. D. The proportion of trajectories pertaining to the four different cases describing the relation of the first stripe-forming MST and the final MST, split by final MST outcome. The four cases are: MST preservation 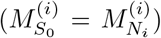, complexification 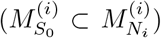, simplification 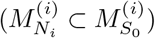 and MST change 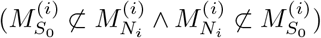.

To classify the mechanistic outcomes of individual trajectories, we identified the minimal subset of regulatory interactions that maintain a basic stripe phenotype, which we term the *minimal stripe topology (MST)*. We determine the MST for the evolved networks at two key time points during evolution: the point of stripe formation, *n* = *S*_0_, and the point at which the target stripe is achieved, *n* = *N*_*i*_, and denote the MSTs at these two time points by 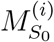 and 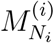 respectively (fig. 1.C and SM, Section 1.6). Using this information we infer a distribution of unique evolutionary outcomes relating to the core mechanisms of stripe formation (fig. 2.B). We identified a total of 732 unique MSTs, with the eight most frequent (MST:1-8) accounting for ~75% of outcomes. The remaining MST outcomes occurred with a mean frequency of 0.034%, and hence this tail represents many different but rare outcomes. We discuss and interpret these rare MSTs and their relationship to MST:1-8 in the Supplementary Text (SM, Section 2).

MST:{1, 2, 5, 6, 7} feature the well-studied feed-forward incoherent type-2 (I2-FFL) mechanism, found in the gap gene network responsible for embryo segmentation in fruit flies^19,47^.

MST:4 is the I3-FFL mechanism^48^, which has not been observed in natural stripe-forming regulatory networks but has been investigated synthetically^13^. MST:{3, 8} feature a negative feedback loop with gene B serving as a controller to determine different responses across inputs, a mechanism which has been shown to enable perfect biochemical adaption^49^. To illustrate that our MST classification captures essential differences in transcriptional dynamics, we solved each of the final networks to steady state and collected concentration profiles at intermediary times^19^ (see SM, Section 1.7). These spatial snapshots of gene expression levels as steady state is approached generate a high-dimensional “fingerprint” for each evolved network and its dynamics (for an examples see Movie 1). A 2-D projection of these dynamical profiles indicates that networks are in general grouped dynamically according to their MST identity (fig. 2.C). This suggests a correspondence between the essential gene interactions for basic stripe formation and the temporal dynamics of gene expression, driven by the complete and (often) more complex network.

In the example trajectory shown in Figure 2.A, the MST associated with the initial and final stripe forming networks are the same 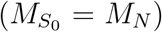. Therefore, after the phenotypic transition to a stripe has been achieved, refinement of the boundaries occurred within the same mechanism. We found that this was often true across our simulations (fig. 2.D). Analogously, developmental patterning typically involves deeply conserved networks, with species specific variations likely achieved through adjustment of interaction strengths^5,7,50^.

We identified 3 additional scenarios related to mechanism preservation across our simulations (fig. 2.D). In some cases, the final mechanism contains additional but necessary interactions whilst maintaining the initial MST interactions 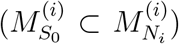, representing “complexification” of the mechanism. Alternatively, the final mechanism represents a simplification of the original 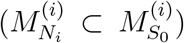, whereby core interactions have been removed without losing the stripe phenotype. Finally in rare cases a fundamental mechanistic shift occurs; at least one core interaction present in both the initial stripe forming network and final network differs in sign 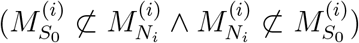. Whilst rare, this latter scenario echoes observations made in bacterial evolution experiments, in which GRNs were selected to have differing responses in a modulating environment^51^. Here, topological innovations were required to overcome conflicting objectives, while changes in regulatory strengths that preserved network topology were sufficient to drive evolution in the absence of conflict.

Overall, the distribution of stripe-forming mechanisms identified in our evolution experiments highlights the many-to-one aspect of the design space of stripe-forming mechanisms as found before^19^. Our analysis also hints towards the differential accessibility of different stripe-forming mechanisms arising from the evolutionary process, possibly dictated by constraints that accumulate during evolution. In what follows we investigate how the mutational process interacts with GRN organization to drive and constrain network evolution and pattern innovation.

### New gene expression boundaries require multiple weight changes per mutant

To understand how mutation interacts with GRN organization and selection for novel patterns, we asked whether specific types of mutations were typically associated with the formation of new gene expression boundaries or the adjustment of existing ones. To this end, we first characterized the phenotypic transition associated with each accepted mutant. Given the mechanistic variation we observed in the final evolved networks (fig. 2.B), we wondered whether there was similar variation in the phenotypic transitions produced through selection. To this end, we characterized the phenotypic transition associated with each accepted mutant.

We reasoned that there are three “routes” to a stripe pattern (fig. 3.A). Starting from the shallow proximal boundary associated with the initial genotype, evolution can first establish a sharp proximal boundary followed by a sharp distal boundary through sequential mutants, do this in reverse, or establish both proximal and distal boundaries simultaneously (colored arrows in fig. 3.A). In between establishing new boundaries, mutation and selection can continue to alter the phenotype, e.g. by adjusting the location or amplitude of existing boundaries (self loops in fig. 3.A). Finally, mutants can be phenotypically neutral, creating no phenotypic changes and no fitness change compared to the resident phenotype.

**Figure 3:**
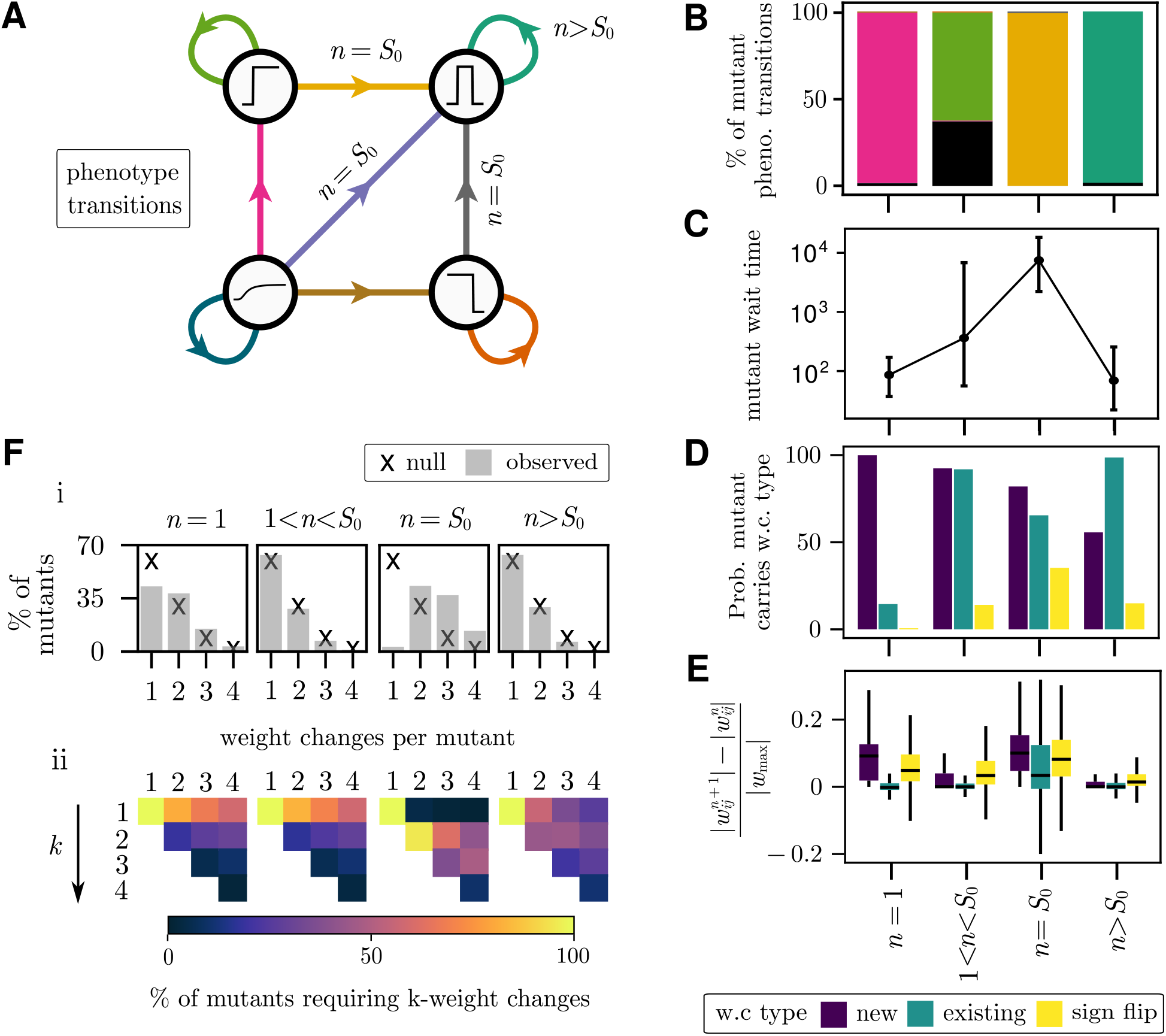
Key components of stripe evolution. A. Phenotypic transitions during evolution. The initial phenotype is that of a shallow proximal-boundary (bottom left). The first accepted mutant at *n* = 1 may lead to a sharp proximal boundary (top left, pink arrow), a stripe (top right, purple arrow), a sharp distal boundary (bottom right, brown arrow) or adjust the existing phenotype without introducing new boundaries (self arrow). Arrows between these states indicate the acceptance of a mutant driving the respective phenotypic changes. Self-arrows indicate accepted mutants that adjust the respective phenotype without adding or removing boundaries. Arrows for transitions that lead to stripe inception are marked by *n* = *S*_0_. B. Percentage of accepted mutants in different evolutionary periods associated with specific phenotype transitions. Colors correspond to arrows in A. The black bar for 1 *< n < S*_0_ corresponds to neutral mutants that do not induce a phenotypic shift or fitness change. C. Median wait times associated with accepted mutants in each evolutionary period (dots). Range bars represent the 25^th^ to 75^th^ percentiles of the wait time distribution. D. Classification of the weight change (w.c.) types carried by accepted mutants in different evolutionary periods as follows: introduction of a new weight (purple), adjustment of existing weight while preserving sign (green) adjustment of existing weight with sign change (yellow). Solid bars indicate the probability of mutants carrying at least one weight change of the type corresponding to the colour. E. The distribution of accepted weight change magnitudes expressed relative to the maximum regulatory effect, defined as 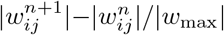, associated with mutants in different evolutionary periods. Color indicates weight change category as in D. F. Percentage of accepted mutants carrying 1-4 weight changes, across different evolutionary periods. Crosses indicate likelihood according to null sampling distribution (pre-selection), bars indicate post selection percentages. For each mutant carrying multiple weight changes we evaluate the minimum number of weight changes, *k*, required to produce the observed fitness gain. This information is visualized in the heatmap, which presents the distribution of required weight changes *k* against the original number carried by the mutants. Yellow along diagonal entries indicates that all weight changes are necessary for the fitness gain.

We found that 98.8% of trajectories introduced the proximal boundary first followed by the distal boundary (Figure 3.B). The vast majority (98.4%), establish a sharp proximal boundary with the first accepted mutant (*n* = 1), followed by adjustment of the newly formed proximal boundary to align it with that of the target stripe (1 *< n < S*_0_). The distal boundary is established at *n* = *S*_0_, with its location then similarly adjusted to match the target pattern during *n > S*_0_. The prominence of this phenotypic path is consistent with the initial network topology, in which the morphogen inhibits output gene *c* (orange) through the intermediary gene *a* (gray) (fig. 1.C). This initial constraint means that creating the proximal boundary simply requires increasing positive feedback on gene *c*. Accordingly, ~ 96% of accepted mutations at *n* = 1 introduce a self-activation interaction on the output gene *c* (orange). The wait times associated with the acceptance of mutants in each evolutionary period indicate that the proximal boundary is typically established quickly (fig. 3.C). In contrast, establishing the distal boundary at *n* = *S*_0_ takes orders of magnitude longer (fig. 3.C). Therefore, despite the degeneracy in the genotype that underlies stripe formation identified in the previous section, phenotypic trajectories are highly stereotyped. These phenotypic trajectories reflect the accessibility of new gene expression boundaries imposed by gene interactions already present in the system at the onset of selection for a stripe-like pattern. Accordingly, altering the initial network configuration impacts the typical sequence of phenotypic transitions (fig.S1).

To understand what type of network changes typically produced each phenotypic transition, we characterized the types of weight changes carried by mutants accepted within each evolutionary stage (fig. 3.D). Mutants accepted in periods 1 *< n < S*_0_ and *n > S*_0_ primarily carry adjustments to existing weights, and weight changes during these periods are typically of small magnitude (fig. 3.D-E). These periods are associated with the movement of existing boundaries, supporting the conclusion that the refinement of existing gene expression boundaries can be achieved through small adjustments to an existing network. In contrast, formation of new boundaries at *n* = 1 and *n* = *S*_0_ typically required the addition of novel regulatory interactions (i.e. new edges in the GRN; fig. 3.D) and was often associated with weight changes of larger magnitude (fig. 3.E). Additionally, distal boundary formation was frequently associated with a change in sign of an existing regulatory interaction, from inhibition to activation or vice versa (fig. 3.D). Therefore our model suggests that generating novel gene expression boundaries typically requires more extensive network re-organization than the adjustment of existing boundaries. These findings are consistent with observations made in the fly embryo, whereby point mutations in existing enhancers led to changes in expression levels that were typically restricted to the WT pattern, whereas new patterning phenotypes emerged through insertion of novel enhancer elements^9^.

To investigate whether the number of weight changes in individual mutants impacts fitness, we next characterized the number of weight changes per accepted mutant in each evolutionary period (fig. 3.F). Pre-selection, the probability of a single weight change, *p*, specifies a null distribution for the number of weight changes per mutant (crosses, fig. 3.F[i] - for derivation see SM, Section 1.2.1). Post-selection this null distribution may be “rebalanced”, representing the number of weight changes carried by accepted mutants, and may be suggestive of a selective bias towards more or less weight changes (bars, fig. 3.F[i]). Multiple weight changes in a single mutant could emerge due to correlations between binding affinities of different TFs to CREs, due to the proximity of their binding sites, cooperative or competitive effects^52–54^. Alternatively, in rare cases, two independent mutations at the sequence level may appear in the same mutant.

For *n* = 1 and *n* = *S*_0_ we noted a bias for more weight changes in accepted mutants in the post-selection distribution compared to the null distribution. The probability that a stripe-forming mutant carries only a single weight change is ~ 3%, compared to the null expectation of ~60% (fig. 3.F[i]). A similar bias, albeit less pronounced, is observed for *n* = 1 (fig. 3.F[i])). Conversely, the number of weight changes carried by mutants in periods 1 *< n < S*_0_ and *n > S*_0_ follow the null distribution. Taken together, these results suggest that the generation of new gene expression boundaries requires interactions between multiple genes in the network to change simultaneously. Inversely, adjustment of existing boundaries is more readily achieved through single weight changes.

To investigate whether our results depend on how rarely two weight changes appear together, we repeated our analysis for smaller values of *p*, reducing the frequency at which multiple weight changes appear in the same mutant. Notably, the post-selection distribution of weight changes per mutant across the different evolutionary periods was invariant for smaller *p*, but the wait times for stripe inception increased as expected given the increasing rarity of the multiple mutation for smaller *p* (fig.S2.A-B). This finding suggests that the capacity to generate multiple weight changes at once may impact the evolvability of GRNs and the spatial patterns they regulate during development. We elaborate on possible constraints at the regulatory level that could therefore either hinder or facilitate pattern evolution in the discussion.

For each mutant carrying multiple weight changes we additionally evaluated the minimum number of weight changes that were strictly necessary to achieve the observed fitness gain (fig. 3.F[ii]). This evaluation reveals 1) when multiple weight changes were strictly necessary for the original fitness gain and 2) when weight changes interact with one another to impact the phenotype. The quantification shown in Figure 3.F[ii] reveals that many of the initial mutants (*n* = 1) carrying multiple weight changes achieve the associated fitness gain using only a single weight change. This is puzzling as the post-selection distribution of the number of weight changes per accepted mutant suggests a selective advantage for multiple weight changes. We reasoned that this selective advantage for multiple weight changes at *n* = 1 can be understood in the context of the minimality of the network at this stage of evolution. Because few interactions are initially present, new interactions are more likely to have no regulatory effect on the output gene and hence be phenotypically neutral. Introducing more weight changes merely increases the likelihood that an interaction between genes with a positive fitness effect is introduced (fig.S3).

Inversely, at *n* = *S*_0_, the increasing number of weight changes carried by accepted mutants is necessary for the associated fitness advantage (fig. 3.F[ii]). Given that multiple weight changes per mutant is a necessary feature for the evolution of new gene expression boundaries in our model, we sought to mechanistically understand how these interactions worked in combination to facilitate evolutionary trajectories. To this end, we investigated the quantitative effects of individual versus combined weight changes on fitness. Epistatic effects naturally arise within gene regulatory networks since by definition multiple genes interact to activate or inhibit other nodes in the network^55,56^. Therefore, placing our analysis in the context of epispastic interactions is a promising framework for our analysis.

### Epistatic effects explain the need for multiple weight-changes

A broad definition of epistasis implies non-additive fitness effects between individual mutations^57^. Epistasis in our model emerges when the sum of the fitness changes associated with individual weight changes does not equal the fitness change induced by the combined weight change mutant. Epistatic effects can be classified into different types. In a 2-weight mutant which is adaptive (i.e. Δℱ *>* 0), sign epistasis refers to a situation where at least one of the changes is deleterious in absence of the other. When both changes are deleterious individually this is known as reciprocal sign epistasis. Magnitude epistasis refers to the case where both individual changes are adaptive but together produce a fitness effect which differs from the sum of their individual contributions, and can be positive or negative^57^ (for quantitative definitions see SM, Section 1.8.2).

To understand the role of these epistatic effects in our evolutionary process, we focused on mutants carrying two weight changes, and characterized their epistasis type across the four key evolutionary periods we have identified. Firstly, we noted that the frequency at which two-weight mutants exhibited epistasis increased as evolution progressed (Figure 4.A). This is an intuitive result, as the addition of more interactions in a network creates dependencies between its connected components. Secondly, the point of stripe inception was strongly correlated with elevated sign and reciprocal sign epistasis, with the vast majority of two-weight mutants at this point in evolution showing some form of epistasis (Figure 4.A).

**Figure 4:**
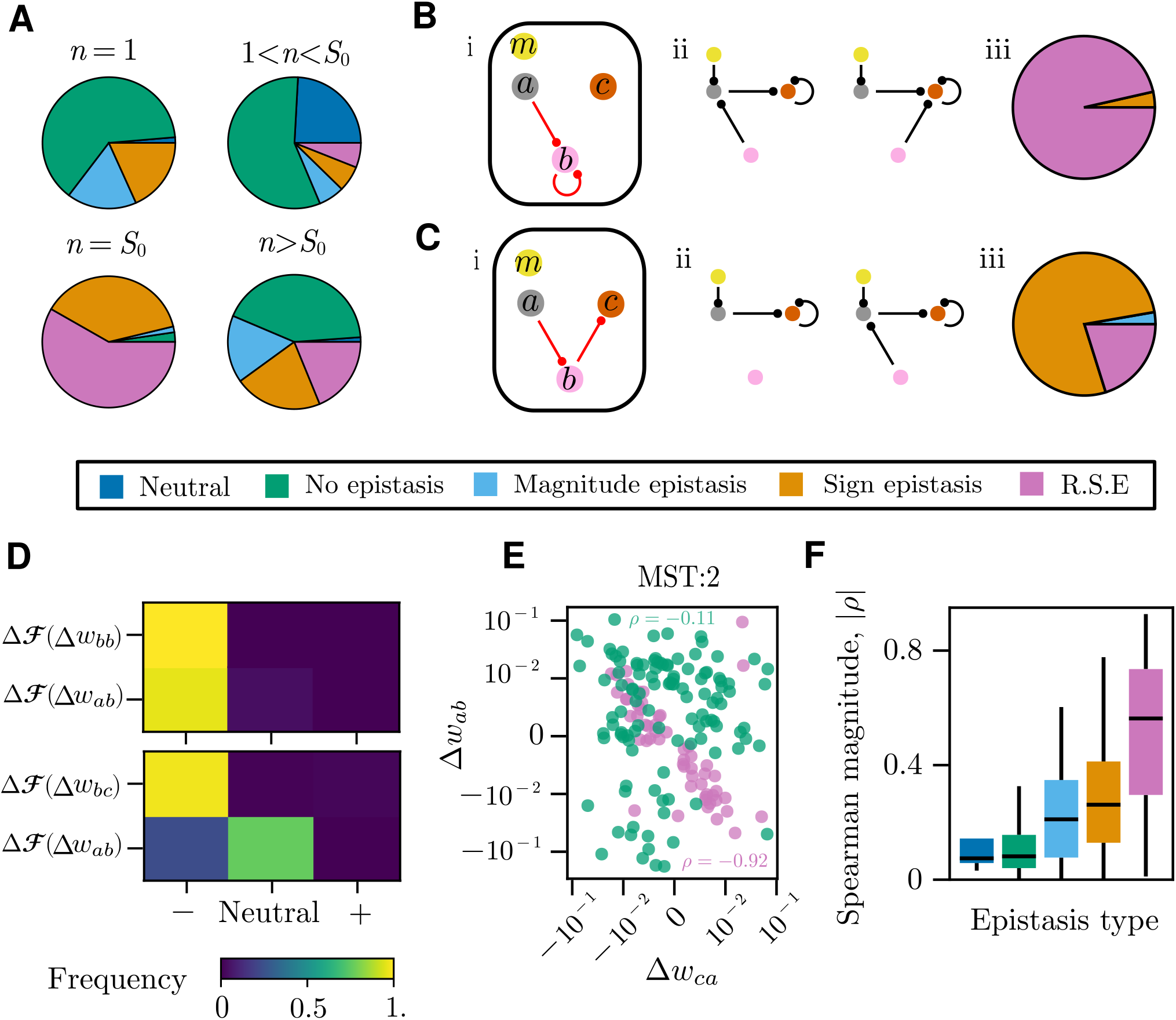
Epistatic effects in 2-weight mutants. A. The distribution of epistatic effect classifications, as defined in the main text and with color coding specified in legend, measured across 2-weight mutants in four different evolutionary stages. B. An example of an mutant at the point of stripe inception, *n* = *S*_0_, with high prevalence of RSE C. An example of an *S*_0_ mutant with high prevalence of sign epistasis. In both panels B and C: (i) The boxed GRN visualization shows the weight changes carried by an example 2-weight mutant and does not show the remainder of the existing network (ii) the top two most frequent network structures prior to mutation (iii) the distribution of measured epistatic effects for this *S*_0_ mutant across our evolutionary simulations. D. The heatmap shows the frequency of fitness effects associated with each individual accepted weight change for the two mutants in panels B[i] and C[i]. Individual weight changes are classified as adaptive (+), neutral, or deleterious (-). For example, instances where ℱ(*w*_*ij*_) and ℱ(*w*_*kl*_) are deleterious would represent a 2 weight mutant with R.S.E. E. Example calculation of weight change correlation. We calculated the weight changes 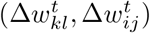 induced by accepted 2-weight mutants, where 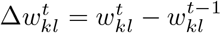. We then measured the correlation *ρ* after grouping 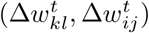 by unique weight identity, MST and epistasis type. F. The distribution of spearman correlation magnitudes measured between pairwise weights, calculated from all 2-weight mutants conditioned on epistasis type.

To understand this, first note that stripe formation in our evolved networks is achieved largely through incoherent feed-forward or negative feedback loops (fig. 2.B). Accordingly, at *n* = *S*_0_ nearly all (99.6%) trajectories carried at least one weight change in the regulatory pathway that connects the gene that is activated by the morphogen (gray) to the stripe forming gene (orange) via interactions with the third gene in the network (pink) (fig.S4). These connections are necessary to form a variant of feed-forward or feedback loop and hence to define the distal boundary.

To investigate why the formation of these mechanisms come hand-in-hand with epistatic effects, we examined the most frequent 2-weight mutants that led to stripe inceptions at n = *S*_0_ (fig.S5.A). Our data indicates that reciprocal sign epistasis frequently arises at *n* = *S*_0_ since gene *b* (pink) has an existing role in regulating the stripe forming gene *c* (orange) prior to stripe formation, either directly or through another gene (e.g fig. 4.B, fig.S6.A). Therefore, any alteration of gene *b* activity must be appropriately compensated by the second mutation, else its individual effect is often deleterious. Sign epistasis at this point emerges when one of the two weight changes is neutral and the other is deleterious when introduced individually. To explain this observation, note that the construction of a feed-forward/feedback loop requires the inclusion of gene *b* (pink). However frequently just prior to stripe inception at *n* = *S*_0_ − 1 gene *b* has no existing regulatory effect on the phenotype (e.g fig. 4.C, fig.S6.B). Therefore, any new regulatory interaction which alters gene *b* without additionally introducing an interaction between gene *b* and the stripe forming gene c (orange) will be neutral. It follows that an increased capacity for accepting neutral mutants may allow trajectories to bypass the constraints imposed by sign epistasis at stripe inception.

To test the effect of neutral evolution, we increased the likelihood of accepting neutral mutants, and restricted the mutational process to generate only single weight changes per mutant. In this high-drift regime mutants that provide no fitness advantage, and so are evolutionarily neutral, can fix at higher probability. Under these conditions, new gene expression boundaries at *n* = 1 and *n* = *S*_0_ were frequently established via the sequential fixation of a single weight change at a time (fig.S7,S8). So whilst a selective advantage exists for mutants which induce multiple regulatory changes simultaneously, neutral mutations and drift may contribute to the evolution of new gene expression boundaries. This occurs when weight changes of little or no effect on fitness accumulate simply by chance but provide a fitness advantage to future single-weight change mutants. In this way our findings suggest that neutral drift may contribute to the evolvability and diversification of GRN-driven patterning.

The overall increase of epistasis as evolution progressed in our model is to be expected, since stripe-forming mechanisms represent an increasingly connected set of regulatory interactions supporting non-linear patterning dynamics. Indeed the correct function of feed forward mechanisms involves precise interactions between participating genes^58^, which can arise through mechanistic requirements such as maintaining counter-balanced gene-gene interactions^59^. We reasoned that this increased mechanistic dependency between genes is likely to induce correlations in the underlying regulatory weight changes post selection. To test if this was true in our model, we calculated the weight changes 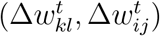 induced by accepted 2-weight mutants, where 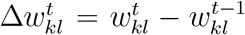. We then measured the correlation *ρ* after grouping 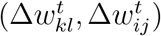 by unique weight identity, MST and epistasis type (fig. 4.E). As predicted, the correlations between weight changes are higher in the presence of epistasis and maximal in mutants that exhibit RSE (fig. 4.F). Furthermore, we found that RSE was particularly enriched when considering weight changes within the MST sub-network (fig.S9). These findings echo previous theoretical work whereby the primary dynamics of biological systems and their evolution depend on a reduced set of system parameters, effectively constraining the possible directions along which evolutionary change can occur^60,61^.

Together these results indicate that epistatic effects strongly constrain evolutionary exploration in patterning evolution as has been demonstrated in other systems^15,62,63^. Therefore, regulatory constraints that induce dependencies between different gene-gene interactions may facilitate or hinder the evolutionary process. The effects of these dependencies on evolutionary paths is likely to depend on on whether the direction of increasing fitness and the induced correlations are orthogonal or parallel.

### Constraints and predictability during GRN evolution

In the context of evolution, epistasis is known to constrain evolutionary trajectories^15^. In particular the presence of reciprocal sign epistasis (RSE) represents “valleys” in the fitness landscape, which are less likely to be crossed through natural selection, mutation and drift^62,63^. Moreover, epistasis has been shown to typically increase as evolution progresses^64^. Whilst epistasis has been identified in patterning networks^13,65^, its emergence in a mechanistic context and its association with specific network changes within an evolutionary process is relatively unexplored. Epistasis also provides an important context for the classic question of historical contingency^66^, in which early chance mutations can restrict evolutionary paths^16,59,67,68^, committing evolution to particular yet highly contingent outcomes.

We therefore wondered to what extent future evolutionary paths in our model become contingent upon the weight changes accepted in intermediate steps. Whilst it is theoretically understood that evolutionary paths in high dimensional genotype spaces become increasingly constrained as evolution progresses,^69^, the mechanistic underpinnings of this phenomena, particularly in the context of complex developmental mechanisms, is not clear^70^. Our framework generates a large number of evolutionary trajectories at high efficiency. Thus, we can obtain data sets that are sufficiently large to investigate the predictability of evolution using statistical methods. Using our large dataset of trajectories, we next aimed to identify mutations that hold a pivotal role in influencing evolutionary outcomes, despite the inherent randomness within the evolutionary process.

We built a predictive model of MST outcome given the regulatory interactions present at any point in an evolutionary trajectory. We set a prediction task that can be described as follows. Given an evolutionary trajectory which has not yet achieved a stripe, predict a set of probabilities representing the likelihood of each possible mechanistic outcome (MST). In other words, given some partial trajectory of genotype evolution 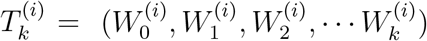 with 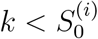, we want to predict a set of probabilities 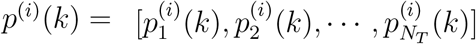 where *N*_*T*_ is the number of MSTs identified and 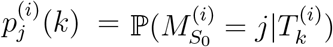 is the probability that the MST associated with the stripe forming mutant at *n* = *S*_0_ is the j-th MST. This is a probabilistic classification task, where the history of the evolutionary process up until the prediction point constitutes the explanatory features, and the initial MST at *n* = *S*_0_ represents the “class outcome”. The learning problem simplifies considerably under the observation that our evolutionary process is Markov, and therefore given a partial trajectory 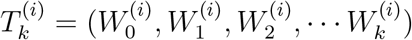, the most recent state 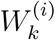 is all that is required to make a prediction about the future. We can then use the trained classifier to yield probabilities for the different MST outcomes given the evolved network at any point in a trajectory. For the purpose of training a probabilistic classifier we limited ourselves to the top 4 MST’s (65.4% of trajectories; fig. 2.B), with an additional “Other” category representing the remaining MST examples (34.6%). Further details of the training procedure can be found in SM, Section 1.9.

An example trajectory and its associated predictions is shown in Figure 5.A. This trajectory consisted of four accepted mutants before stripe inception at *n* = *S*_0_ = 4. The initial prediction *p*^(*i*)^(0) simply reflects the proportion of each MST observed in the train-set – the null prediction learned by the model in absence of any information on the further evolution of the network. Following the first accepted mutant a new prediction, *p*^(*i*)^(1), indicates that the network changes have increased the likelihood that MST:1 will be adopted at the point of stripe formation. Following the second accepted mutant MST:4 becomes the most likely outcome, and forms a correct prediction which persists until *S*_0_. We describe a mutant which changes the MST outcome prediction as branching (see SM, Section 1.9). Therefore in the above example, the second mutant was branching. In this way our model suggests that a single mutant can drastically change the likely evolved stripe-forming mechanism. These types of mutants are important since they represent the incorporation of regulatory constraints which change the likely path of evolution. Therefore, this quantitative framework allows network changes driving specific mechanistic outcomes to be identified automatically at a large scale.

**Figure 5:**
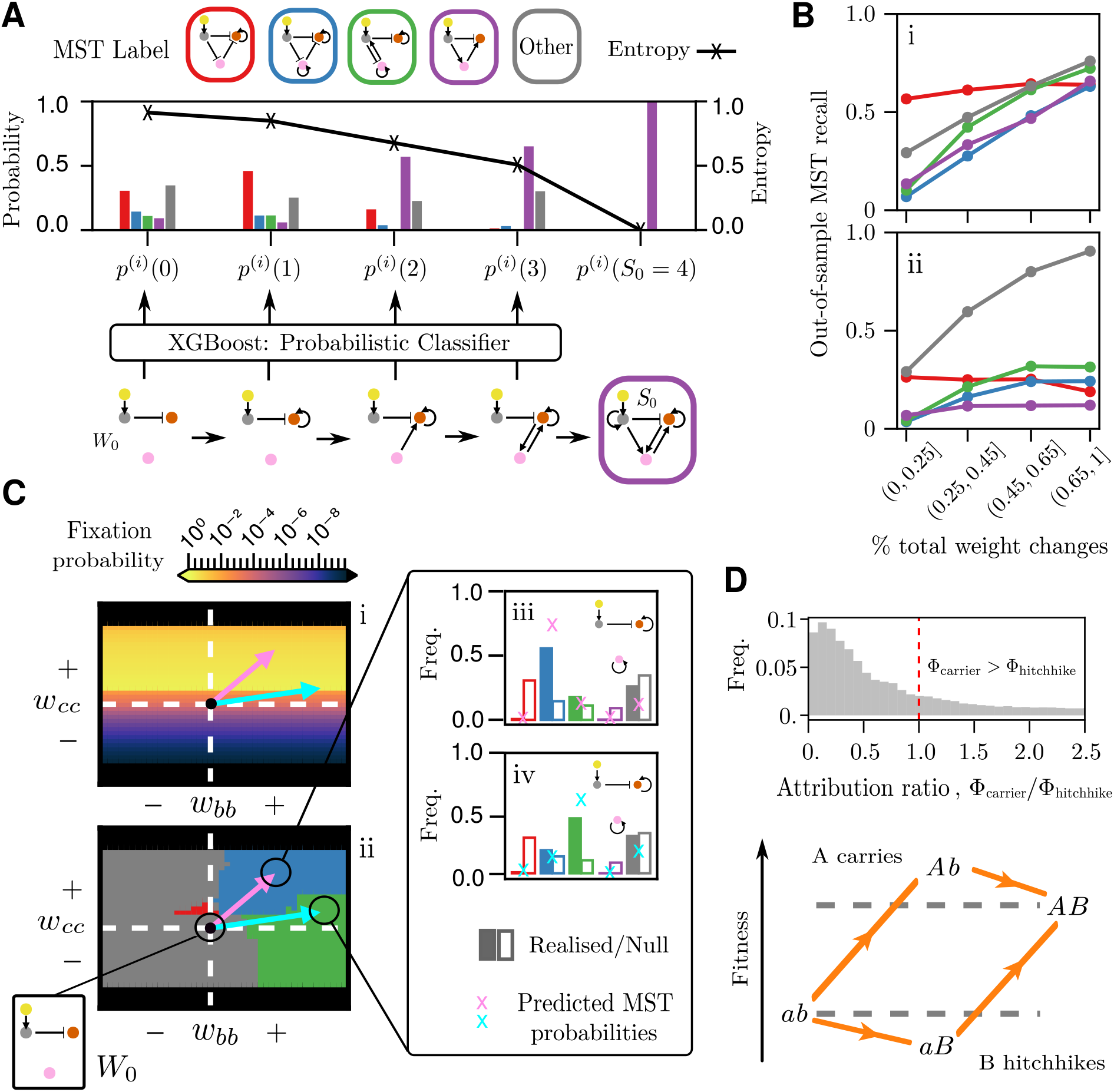
Predictability of stripe evolution. A. An example trajectory and the MST prediction probabilities for intermediate mutants. Colors (Red,Blue,Green and Purple) represent MST labels and correspond to color coding used in Figure 2, with gray representing the “other” MST category. A sequence of network configurations corresponding to fixed mutants in an example trajectory *i* is shown, with the MST realized at *S*_0_ shown highlighted in purple, corresponding to MST:4. For each network 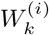 in the *i*−th trajectory we produce a probabilistic MST prediction *p*^(*i*)^(*k*) using a trained XGBoost classifier. Here 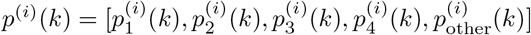 where 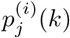 is the probability that the *j*-th MST is realized at *S*_0_ given the *k*-th mutant. Coloured bars indicate the individual MST prediction probabilities 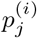. Black line indicates entropy associated with the prediction probabilities *p*^(*i*)^(*k*). B. The recall for each MST outcome evaluated on out-of sample networks at different stages of evolution, evaluated using the complete predictive model (i) and the topology only predictive model (ii). Networks were first binned according to the number of weight edits they had accumulated as a percentage of the total accumulated at *S*_0_. The bins were chosen so that the number of networks in each bin was approximately equal. Next, the recall for each MST outcome was evaluated for the networks in each bin. Recall is the % of correct predictions for the outcome. A prediction is deemed correct if it matches the MST outcome and furthermore, persists from the point of prediction to *S*_0_ - for more details see SM. C. 2-dimensional fitness landscapes associated with pairwise weight changes carried by the (*n* = 1) mutant (i) with the associated MST predictions (ii). Pink and cyan arrows indicate weight changes associated with selected branching mutations. Bar charts (iii) and (iv) represent the MST outcome distributions associated with pink and cyan branching mutations. Filled bars indicate the distribution induced by the branching mutations, whereas non-filled bars represent the null distribution 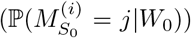. Scatter crosses indicate probability of each MST outcome given by predictive model for the branching mutants. D. Histogram showing the distribution of prediction attribution ratios Φ_carrier_*/*Φ_carrier_ associated with mutants carrying hitchhiking weight changes. Red dashed line indicates equal attribution between hitchhiking and carrier weight changes. The schematic shows that hitchhiking weight changes (*b* → *B*) are those which are deleterious individually, but adaptive when fix with a carrier mutation (*a* → *A*).

In general we were able to predict final MST outcome prior to *S*_0_, with a good out-of-sample predictive performance given relatively few network modifications (fig. 5.B, SM, Fig.M1). Interestingly, the predictive performance of the model deteriorated significantly when only topological information was considered by the classifier (fig. 5.B[i] versus [ii]), indicating that network topology alone is not sufficient for determining the likely path of evolution and that the magnitude of the interactions between genes matter.

We quantified the statistical entropy, *H*(·), associated with the predicted probabilities *p*^(*i*)^(*k*) changes as mutants fix (Figure 5.A). Interpreting the entropy as a measure of uncertainty inherent to the MST outcome, this example highlights the intuitive notion that uncertainty decreases as evolution progresses. We found this to be a general principle, with mutants and their corresponding network changes most often decreasing uncertainty in eventual MST outcome (fig.S10.A-B). This is in agreement with previous work showing that the space of possible evolutionary paths that a trait can take tends to narrow during its evolutionary history^70,71^. Despite this been an intuitive and established result for abstract genotype-phenotype models^69^, our analysis identifies possible mechanistic origins for these constraints in the context of developmental GRNs.

We then wondered whether we could steer evolution towards a particular MST outcome by “engineering” the initial mutant. We used the predictive model to identify initial (*n* = 1) mutants that were branching towards a specific MST outcome and identified the branching mutant producing the largest negative entropy change, implying a highly certain mechanistic prediction. To aid tractability, we focused on mutants carrying 2 weight changes relative to the initial network. The mutants found through this procedure for MST:2 and MST:3 both induce the same change in fitness and hence have an equal probability of fixation (fig. 5.C[i]). Interestingly, these mutants induce the same topological change, however differences in the weight values result in alternative mechanistic predictions (fig. 5.C[ii]). We then imposed the identified mutations at *n* = 1 and simulated 5000 independent trajectories from these new initial conditions. In both instances evolution was steered towards the predicted mechanism (fig. 5.C[iii-iv]). Similar results were observed for mutations predicted to steer evolution towards other MSTs (fig.S11). These findings suggest that regulatory interactions incorporated early in network evolution could significantly bias evolution towards a specific mechanistic outcome.

Recall that multiple weight changes were often selected for together at the fist accepted mutant (*n* = 1), not all of which were strictly necessary (Fig. 3 F). Each adaptive weight change at *n* = 1 therefore likely fixed with a non-adaptive weight change, reminiscent of genetic hitchhiking whereby neutral or slightly deleterious mutations appear together with a beneficial mutation and increase in frequency as a result^72^. Neutral or nearly neutral network changes may provide a reservoir of cryptic genetic variation for subsequent evolution^73^, which enables paths to new adaptive genotypes and creates historical contingency in protein evolution^74^. We explore this idea in the context of regulatory evolution by using our predictive model to identify which weight changes carried by a mutant contributes most strongly to its mechanistic prediction.

For each weight change we produce a prediction attribution, Φ, which quantifies to what extent an individual weight change shifts the prediction for the MST outcome of evolution (see SM, Section 1.9.5). To then identify the relative importance of the hitchhiking versus adaptive weight change in each mutant, we calculate the ratio Φ_carrier_*/*Φ_hitchhike_. In the majority of cases, the hitchhiking weight change is far more instrumental in determining the mechanistic outcome than the weight change associated with the carrier (fig. 5.E). This suggests that regulatory networks may accumulate mutations which are selectively-neutral but nonetheless significantly alter the course of subsequent evolution. A similar conclusion has been reached experimentally in a transcription network that controls the response to mating pheromone in yeast^75^, where the gain of a neutral regulatory interaction permitted changes in the overlapping circuit responsible for pheromone induction.

## Discussion

In this work we have examined how developmental mechanisms interact with evolutionary processes. By placing GRNs in an evolutionary context we have investigated how gene network organization, mutation and natural selection together determine evolutionary outcomes. Changes at the regulatory level is a fundamental avenue through which developmental patterns diversify during evolution^2–8^. Although our analysis focuses on evolving stripes, forming and adjusting gene expression boundaries is fundamental to developmental patterning and its evolution. Therefore, our findings and conceptual framework may offer general insights towards a mechanistic understanding of how regulatory networks that drive spatial patterns emerge and diversify.

By framing the evolution of a stripe in gene expression levels as successively establishing and adjusting sharp gene expression boundaries, we identified network modifications commonly linked with specific phenotypic innovations. Our findings suggest that adjustments of existing boundaries typically occur through smaller changes in the strength of existing gene interactions. Inversely, novel boundaries are achieved through mutations that impact multiple gene interactions and have a larger effect on network organization. These findings are consistent with recent mutagenic studies in *Drosophila* embryos^9^ where point mutations on native enhancers rarely resulted in expression outside of the native pattern, whereas inserting *de novo* enhancer elements often drove novel patterns. Our model offers a mechanistic theoretical framework to interpret these experimental observations. Point mutations in binding sites or TF coding regions typically induce smaller changes in transcription rates^39,76^, whereas larger changes can arise through whole enhancer duplication^77^. Alternatively, a novel enhancer element may induce a new regulatory interaction into the network^78^. Therefore, although we do not explicitly model gene regulation at the sequence level, our model is able to capture key aspects of GRN response to mutation and corresponding phenotypic shifts.

The capacity to generate changes in multiple gene interactions within the same mutant was crucial to forming and adjusting gene expression boundaries in our model. In our analysis, we have placed this observation in the context of functional epistasis, that is known to characterize patterning GRNs^13,65^. Accordingly, epistatic effects were prominent throughout evolutionary simulations. Specifically, sign and reciprocal sign epistasis, whereby two changes in gene interactions are adaptive only when they appear in the same mutant, were key to the formation of new gene expression boundaries and boundary adjustment at later stages of evolution when the core network becomes established. The ubiquitous presence of these epistatic effects across more than 10^5^ independent evolution trajectories indicates that they may be a general property of GRN-driven pattern evolution. Given this prevalence of epistasis in our analysis, we speculate on its possible implications for the evolvability of GRN-driven patterning mechanisms.

A pervasiveness of sign and reciprocal sign epistasis during GRN evolution implies that the capacity to generate adaptive mutants depends on the likelihood of generating single mutants where multiple gene interactions change. Therefore the evolvability of GRN-driven patterning, broadly defined as the ease with which new phenotypes are accessed, depends on the genetic constraints which determine how likely multiple gene interactions are to change together in single mutants. Indeed convergence rates in our simulations correlated with the likelihood of generating multiple weight changes at once. A key question then becomes: how does the molecular basis of transcription and regulation interact with mutation to re-wire regulatory networks?

Experimental work has characterized the evolutionary potential of mutations at the TF level for network rewiring^53^ and the role of enhancers in introducing novel genes into a regulatory network^78^. In our model, specific combinations of gene interactions were selected for together and in a highly correlated manner. Could genetic architecture itself facilitate the correlations that are necessary to resolve these constraints and hence facilitate the evolution of patterning GRNs? For example, theoretical work in a class of genetic algorithms suggests that correlations in system parameters may enable mutations to align optimally with the contours of a fitness landscape, resulting in faster convergence towards fitness optima^79^. In a regulatory network, these type of correlated changes could arise through mutations in enhancer binding sites with cooperative or competitive effects, or when binding sites of independent TFs are in proximity of one another^52,54^. Our work provides ground for studying these effects and their potential role in the evolution of patterning mechanisms. Advances in synthetic biology are beginning to shed light on how fragments on individual enhancers drive TF expression by engineering synthetic CREs and screening for their function^80^. This approach could be combined with our framework in the future to elucidate how the logical rules that underlie enhancer signaling impact GRN-driven spatial pattern evolution. Furthermore, insights from theoretical frameworks that study regulatory evolution by modeling mutation at the sequence level^81^ could be incorporated in our framework to explore their consequences for patterning evolution.

We showed that in principle neutral drift could bypass epistatic constraints by allowing the accumulation of multiple weight changes in the same individual sequentially. Therefore in spite of the strong preference for multiple weight changes, we saw successful stripe evolution using only single weight changes given an increased propensity for neutral weight changes to fix. In this way, our work provides a mechanistic argument for the importance of neutral adaption in the evolution of regulatory networks^82^. The epistatic effects identified in the evolution of stripe forming mechanisms imply that the ability for neutral adaption to circumnavigate epistatic constraints could be a general property of evolution on patterning fitness landscapes. This may arise due to the inherent degeneracy in network topology for GRN-mediated patterning, whereby multiple network topologies can drive the same spatial pattern,^19^ and the presence of so-called neutral networks, which have been shown using synthetic networks to enable new phenotypes to be accessed through mechanisms such as drift^12^.

The dependence of phenotypic accessibility on GRN topology and the mutational process alludes to historical contingency as an important aspect of GRN evolution, so that network changes accumulated early may influence subsequent evolution^13,18^. We have built on this idea by investigating how the evolution of specific stripe forming mechanisms becomes contingent upon early network organization. By developing a predictive framework we identified “branching mutations” early in evolution, which we were able to show significantly altered the likely mechanistic outcome. This suggests that degeneracy in patterning mechanisms, in combination with the constraints imposed by the existing network on subsequent mutations drives historical contingency. By employing machine learning techniques in combination with a large dataset of evolutionary trajectories we present a novel method to investigate the importance of historical contingency, echoing existing probabilistic characterizations of historical contingency^83^. In particular we demonstrated how explainability techniques^84^ offers a principled way to identify the specific motifs which introduce constraints in a network evolution setting. Furthermore, the identification of branching mutations using our framework could guide synthetic GRN evolution assays in the future.

In conclusion, we have developed a high-throughput and tractable computational approach for GRN-driven developmental patterning that we have used to probe general principles of the evolution of development. Our framework can be adapted in the future to integrate new information from synthetic engineering assays^80^ and simulations at the sequence level^81^ that probe the constraints that regulatory architecture introduces to the evolution of GRN-driven developmental patterning. In addition, the framework we have developed can be adapted to investigate other phenotypic transitions or more complex networks. In this way we hope that our findings will stimulate further research, driving advancements in our understanding of developmental pattern evolution.

## Acknowledgments

We would like to thank Amy Bowen, James Briscoe, Jake Cornwall-Scoones, Leonard Dekens, James DiFrisco, Kabir Hussain, Taniya Mandal and Lewis Mosby for discussions and feedback on this work. This work was supported by the Francis Crick Institute, which receives its core funding from Cancer Research UK, the UK Medical Research Council, and Wellcome Trust.

## Notes

### Competing Interest Statement

The authors have declared no competing interest.

### Summary of Updates

Text clarifications and focus, methods details, expansion on mutational effects.

